# The HexMaze: A previous knowledge and schema task for mice

**DOI:** 10.1101/441048

**Authors:** Alejandra Alonso, Levan Bokeria, Jacqueline van der Meij, Anumita Samanta, Ronny Eichler, Patrick Spooner, Irene Navarro Lobato, Lisa Genzel

## Abstract

New information is rarely learned in isolation, instead most of what we experience can be incorporated into or uses previous knowledge networks in some form. However, most rodent laboratory tasks assume the animal to be naïve with no previous experience influencing the results. Previous knowledge in form of a schema can facilitate knowledge acquisition and accelerate systems consolidation: memories become more rapidly hippocampal independent and instead rely more on the prefrontal cortex. Here, we developed a new spatial navigation task where food locations are learned in a large, gangway maze – the HexMaze. Analysing performance across sessions as well as on specific trials, we can show simple memory effects as well as multiple effects of previous knowledge accelerating both online learning and performance increases over offline periods. Importantly, we are the first to show that schema build-up is dependent on how much time passes, not how often the animal is trained.

## Introduction

After infancy, we rarely acquire new information in isolation, instead most of what we learn throughout our lives can be associated with previous knowledge. For example, Harlow (1949) described learning sets as the “learning to efficiently learn” process of generalizing previous experience in a class of problems to new problems of the same class. While most tasks in human behavioural research are based on and embedded in familiar efforts and environments, rodents tend to be naïve to the behavioural tasks and can draw only little benefit from previous experience. This causes one of the largest discrepancies between human and rodent research.

This distinction is critical since the rate of learning and consolidation as well as neural underpinnings and relevant brain areas can show a shift in the presence of previous knowledge (Genzel & Wixted, 2017; Squire et al., 2015; van Kesteren et al., 2012; Wang & Morris, 2009). In human research, the previous knowledge effect has been long established (Bartlett, 1932) but it was not introduced to rodent research until the seminal study of paired-associates task introducing the schema effect on systems consolidation (shift to hippocampal independency) in rats (Tse et al., 2007). During the task, rats initially learn six flavour-location associations: they receive a flavoured pellet in the start box and learn that more of the same flavoured pellets can be found in one specific sand well within the map. After learning six flavour-location pairs over nine weeks creating a mental map or ‘schema’, this map can be updated with new flavour-location pairs. In a sequence of papers, it was shown that previous knowledge accelerates learning to a one-trial event as well as the rate of systems consolidation from weeks to days (Bethus et al., 2010; Tse et al., 2007). Further, in addition to the hippocampus, the medial prefrontal cortex needs to be active during encoding for memories to last (Tse et al., 2011; Wang et al., 2012). The involvement of the medial prefrontal cortex as a structure for the schema effect was then later confirmed in humans (Ghosh & Gilboa, 2014; van Buuren et al., 2014; van Kesteren, Fernandez, et al., 2010; van Kesteren, Rijpkema, et al., 2010).

Later on, Richards et al. (2014) used the watermaze to investigate previous knowledge effects in mice as well as to test if systems consolidation leads to a more gist-like quality in memory instead of individual item representation (Ghosh & Gilboa, 2014; Gilboa & Marlatte, 2017; Hardt & Nadel, 2017), as this was not distinguishable in the paired-associates schema task. They trained mice on daily switching platform locations, which were drawn from a pre-determined statistical map in space, and could show that search patterns in the watermaze faithfully represented the statistical distribution of platform locations. This effect was pronounced even more so at 30 days than at one day after training. Further, they could show that the prefrontal cortex was necessary for updating behavioural search patterns to conflicting new information (a platform incongruent with the previous pattern) if the event occurred 30 days after learning the initial pattern. This was not the case one day after learning. However, no comparison was made to animals without previous knowledge. The Object Space Task (Genzel et al., 2019) uses a similar approach testing the extraction of statistical spatial patterns but in the context of an object exploration task. Laboratory rodents have the natural tendency to explore objects that are more novel, for example if they are moved to a new location since the last encounter. In the Object Space Task objects are presented in predetermined configurations across multiple sample trials to test if animals can accumulate statistical patterns across events. The key condition testing memory abstraction has always one object in one location, with a second, identical object present in one out of three other locations. The final sample and test trial always have the exact same configuration. Interestingly, during the test trial the animals show a preference for the less often shown location (across the previous sample trials) despite the fact that the final sample trial had the same configuration and no change occurred. This shows that both rats and mice are guided by their cumulative experiences across multiple sample trials in their object exploration in a test trial and not by the last sample trial. This once again emphasizes that previous knowledge will affect how an experience, e.g. an interference or additional sample trial, will affect subsequent performance.

In the present study, we aimed at developing a new behavioural task in which we can investigate the role of previous knowledge on new memory acquisition and consolidation across different time-points in training. In order to achieve this, it is important that during both initial build-up of the knowledge network (i.e. schema) as well as later updates of this schema, the difficulty of the task and thereby cognitive load remains the same. Thus, we chose to train mice in a large environment to navigate to a single goal location. We expect to see different types of previous knowledge effects on the performance of the mice, reflected in the length of their navigational paths: learning the general task principles (static food location and allocentric navigation from different starting positions), enhancing memory encoding (increased performance on the second up to the last trial of a session) and enhancing memory consolidation (increased long-term memory and performance on the first trial of each session). To test how quickly new information can be incorporated into this schema (here a map), we changed the goal locations every few sessions. In the HexMaze task, the spatial map of the environment (hexagonal layout, intra- and extra-maze cues) is the schema that should then be – once established – flexibly updated for reward locations or changes within the environment, e.g. the blockage of routes with barriers (Arbib, 1997; Behrens et al., 2018).

We could show that mice need 12 weeks to initially build up a spatial map of this environment, which functions as a schema and facilitates rapid long-term consolidation during later updates. Importantly, this schema build-up is dependent on how much time passes (weeks), not how often the animal is trained (training days). In addition to the enhancement of long-term memory after schema acquisition, we can distinguish a simple memory effect reflected by better performance across the first couple sessions of the first goal location. Furthermore, an initial learning set effect after two weeks of training is seen in the first goal location switch as well as a late learning set effect after 12 weeks of training. This initial learning set effect is not expressed in the first trial of a session (long-term memory) but does facilitate the increase of overall session performance. Finally, focussing on later learning after 12 weeks, we could show that the degree of overlap with previous knowledge influences navigational performance on the first session of a change, i.e. how quickly new information could be incorporated online. Thus, the HexMaze task allows to distinguish four effects of previous knowledge on memory, ranging from learning set to rapid consolidation and within-session updating.

## Results

### The HexMaze

The Hexmaze is arranged as six regular densely packed hexagons, forming twelve two-way and twelve three-way choice points (nodes) 36.3 cm apart, in total spanning 2 m x 1.9 m (Fig. 1A). Gangways between nodes were 10 cm wide and flanked by either 7.5 cm or 15 cm tall walls. Maze floor and walls were white and opaque, with local and global cues applied in and around the maze to enable easy spatial differentiation and good spatial orientation; overall leading to a complex, integrated maze. To measure performance in this maze, we divided the taken path of each trial by the shortest possible path (Fig. 1B, comparison of different performance parameters in Fig. S1). To eliminate the resulting skewness (skewness 3.33) we used the log of the normalized path (skewness 0.72). Each session lasted 30 min per animal, resulting in 25-35 trials per session (Fig. S3) with each trial starting from a different location within the maze (Fig. 1C). Evaluation of the performances of only the first trials of the sessions measures long term memory performance and during critical sessions, e.g. the second session of a new goal location (GL), this first trial was used as a probe trial where the food reward was not present for the initial 60 s to control for olfactory cues. In contrast to the first-trial evaluation for long-term memory, looking at the performance over all trials gives a measure of the overall working memory and navigational performance within the environment.

**Fig. 1.**
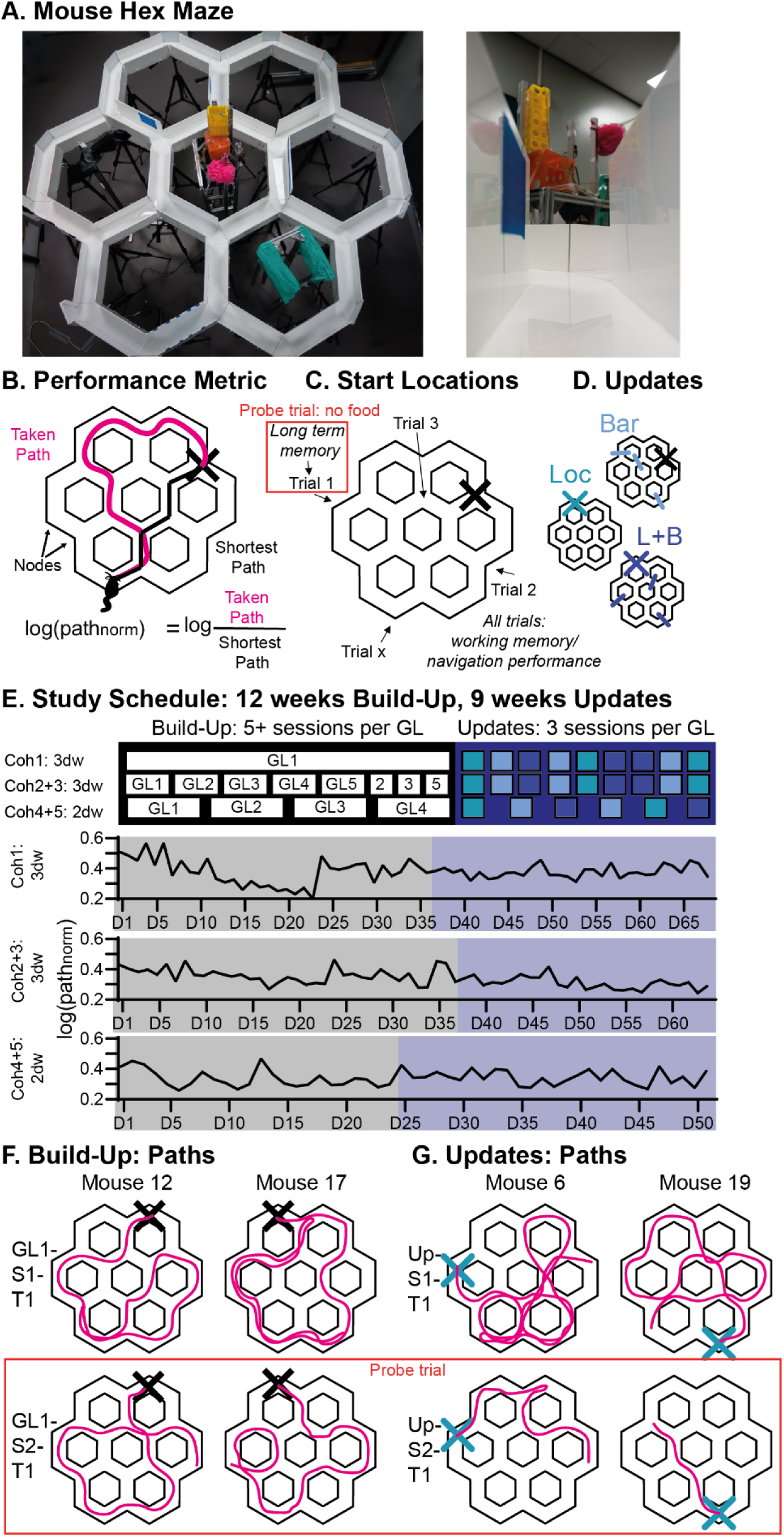
The Hex Maze: **A.** Shows the maze with intra- and extra maze cues (left) and the maze from the view of the mouse (right, see also supplemental video 1) **B**. The main performance metric is the log normalized path (path_norm_) with the length of the paths taken by the animal divided by the shortest possible path to the goal location (GL, indicated by the X). **C.** During training, animals started each trial from a different location and had to navigate to a fixed GL. A first trial measures long-term memory performance and was utilized as a probe trial on critical sessions (no food present). Performance on all trials of the session measure general working memory/navigational performance in the known environment **D**. After the animals had acquired the general maze knowledge during the Build-Up, Updates were performed with inclusion of new barriers (Bar), new goal locations (Loc), or the inclusion of both (L+B). **E.** Shows the general training schedule for all animals during the whole experiment. Animals were trained to one GL in a given session. Cohort 1 (coh 1) only had one GL during the Build-Up and this cohort was used to pilot general principles as maze size, food deprivation etc. (see SF1 and 2). In contrast, for coh 2-5 the GL was kept constant for seven sessions of the first GL (GL1), then five or six sessions for GL2, and five or seven sessions each for GL3-5. For coh 2 and 3, three of the initial five locations were repeated with each three sessions. Finally, for all cohorts, each Update contained three sessions. The sequence of the Update types was counterbalanced across animals. Each Update type was repeated 2-3 times. Throughout all phases the first trial of the second session and during Build-Up first trial of the fourth, fifth, or sixth session were utilized as probe trials. Coh 1-3 were trained three days a week (3dw), coh 4+5 two days a week (2dw). Example paths of the Build-Up and Updates are shown in **F**. and **G**. respectively (Supplemental videos 2-9).

Animals went through two phases of training: Build-Up and Updates. In the Build-Up the animals should create a cognitive map of the maze environment; in contrast, during Updates, stable performance is achieved and they should be simply updating the cognitive map. These two phases also differed in the frequency of GL switches: during Build-Up, the GL remained stable for five and more sessions, while during Updates a change occurred every three sessions (see also below). Different Update types were performed: including barriers in the environment (Bar), changing the goal location (Loc) and doing both (L+B, Fig. 1D).

Five cohorts (coh 1-5) of four animals each were trained in the maze (Fig. 1E). Coh 1-3 were trained three times a week (3dw) while coh 4+5 were trained two times a week (2dw). Coh 1 was trained on only one GL during the Build-Up and was used to pilot the maze size (Days 1 to 23 on a smaller maze, Fig. S2), food deprivation (started Day 12, Fig. S2) and other experimental factors. This data (see Fig. 1E) is not included in the later analysis, instead only the Updates of coh 1 was used (Fig. 4D and E). For coh 2-5, the GL was switched during the Build-Up every 5-7 sessions (GL1: 7 sessions, GL2: 5/6 sessions, GL3-5: 5/7 sessions) to test when rapid updating could occur. Faster switches were initially avoided, to help shape the animal’s behaviour. In the first trial of the day animals would not find food at the last presented location for both the first session of a new GL as well as probe trial days (e.g. always the second session of a new GL); thus these sessions were interleaved with normal training sessions with food present in the first trial at the last known location to avoid the animals learning the rule that food is initially not provided.

**Fig. 4.**
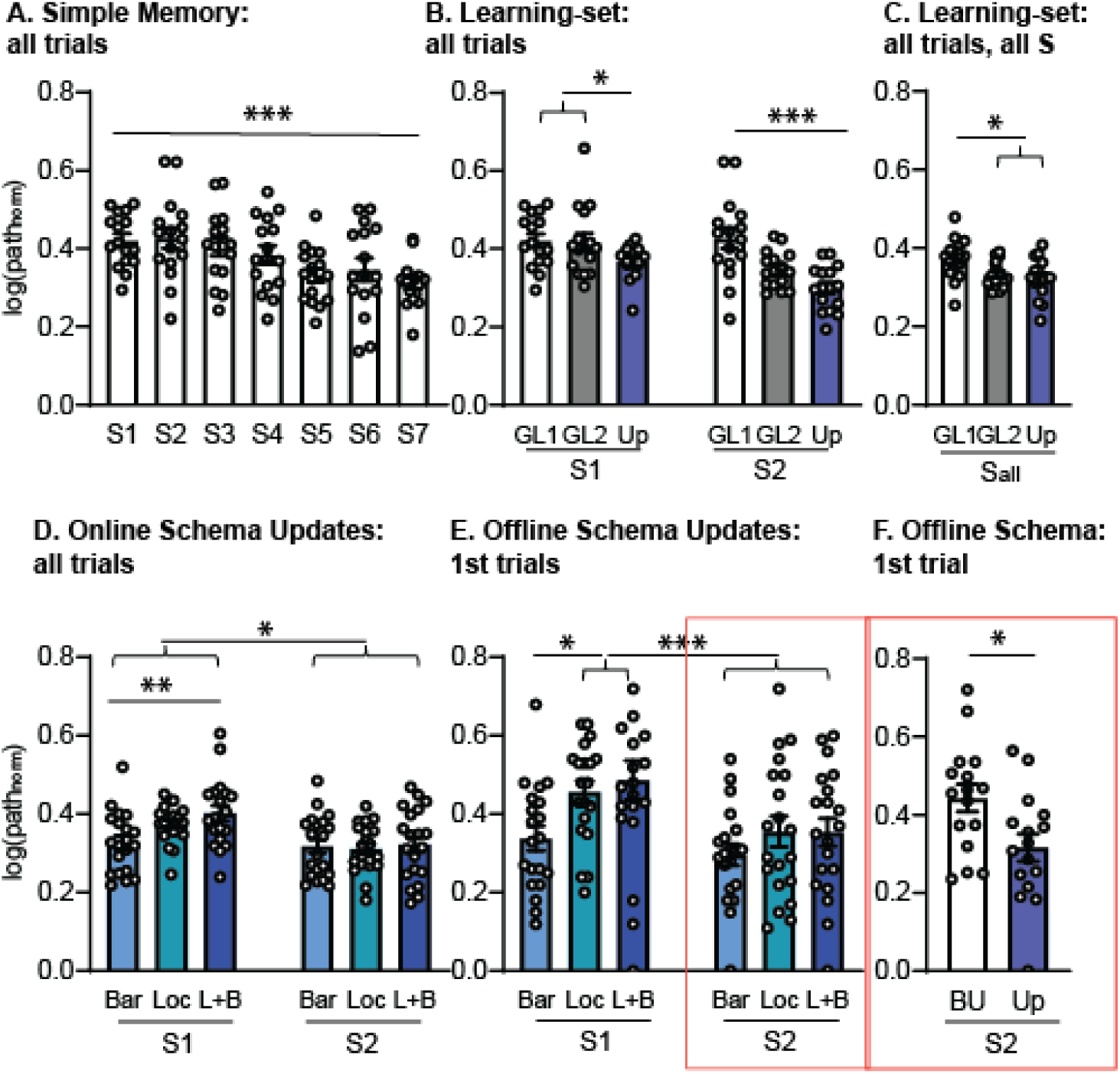
Previous Knowledge effects: **A.** Plotted is the whole session performance for the first GL during the Build-up. The significant session effect reveals a performance increase dependent on experience indicating a more efficient working memory/navigational performance. **B.** Plots the performance for the first two sessions of the first two GLs during the Build-Up, as well as Updates (averaged for all types). Already for the second GL (three weeks since training start) a significant increase in performance (decrease of path length) is seen in the second session in comparison to the first session. This overnight (offline) performance increase is comparable to the increase found after seven sessions for the first GL. This may represent a more efficient consolidation and updating effect but is only expressed in the whole session average (not long-term memory present in the first trial, see Fig. 2 and 3). During the Updates, this performance increase is already visible in the first session with additional offline gains found in the second session. This three-step performance gain is reminiscent of a learning-set effect (Harlow, 1949). **C.** Considering all sessions, we find that animals already reach overall plateau performance by the second GL. **D.** Zooming in on the performance during the first and second session during the Updates, another previous knowledge effect is revealed across the different Update types. The barrier (Bar), goal location (Loc) and combined updates (L+B) differed in their overlap of previous knowledge (or need for updating that knowledge) which influenced how well they performed (all-trial) in the first session. **E**. Shows the same effect but now for only the first trials. Only in the presence of a goal switch did performance in the first session decrease. However, by the second session this performance difference was gone, revealing that one session is sufficient for the memory update. Finally, **F.** depicts the performance of only the first trial of the second session during the Build-up and Updates (only Loc and L+B) where long-term memory (2-3 d) after one session learning to a new GL improves from Build-up to Updates. Thus, it seems once a schema is established, only one session training leads to better long-term memory performance (as also seen in E). Orange boxes indicate that the trial was utilized as a probe trial meaning food was not present for the initial 60 s. The single asterisk stands for p<0.05, the double asterisk for p<0.01 and the triple asterisk for p<0.001.

After 12 weeks of Build-Up, all cohorts were tested in the Updates, where a change (given by the different Update types) was introduced every three sessions. The sequence of the different Update types (Loc, Bar, L+B) was counterbalanced across repetition and cohorts. Further, the GLs were also counterbalanced across animals within a cohort as well as across cohorts. To ensure that the identity of individual GLs did not account for learning effects over time, the sequence was reversed between cohorts, e.g. GL1 of the first animal in coh 2 would be GL5 of the first animal in coh 3.

Overall performance for each cohort across time can be seen in Fig. 1E. Different learning effects were found as highlighted in individual paths (Fig. 1F and G, supplemental videos 2-9): on the first trial of the first training day of the Build-Up, the animals show random movement through the maze and just by chance find the GL (Fig. 1F, supplemental video 2, 4). On the next day at the first trial some but not all animals already show more goal directed behaviour (Fig. 1F, supplemental video 3, 5). In contrast, during the Updates on the first trial of a new GL the animals are more likely to show efficient foraging strategies (efficient exploration of the known environment of the maze, Fig. 1G supplemental video 6, 8) in contrast to random exploration during the initial Build-Up; the difference is notable by animals e.g. revisiting nodes less within the maze. And in the succeeding session of the Updates most animals showed goal-oriented navigation to the reward location already on the first trial (Fig. 1G, supplemental video 7, 9).

### Building and updating the map

To formally investigate the effects seen in the individual paths, we analysed group level performance in more detail. Coh 2+3 (total n=8) were trained Monday/Wednesday/Friday (example study schedule Fig. 2A) and during the Build-Up showed a significant improvement in navigation to the GL (all-trials) across sessions as well as across GLs (GL1-5, Session F_4,28_=6.2 p=0.001, GL F_4,28_=3.3 p=0.026, interaction F16,112=1.4 p=0.15). For both session and GL, the linear contrast was significant (Session p=0.027, GL p=0.043. Fig. 2B). In the Updates, the animals overall performed better than in the Build-Up (F_1,7_=8.2 p=0.024), and continued to show a significant improvement of performance over the three sessions (Session F_2,14_=12.9 p=0.001, linear contrast p=0.005). Additionally, there was an effect of Update type (Bar, Loc, L+B) as well as a type X session interaction: in contrast to the Update types with location changes, animals already performed well in session 1 of the barrier updates (type F_2,14_=3.5 p=0.058 with linear contrast across Bar, Loc, L+B p=0.027, interaction F_4,28_=2.6 p=0.059, orthogonal comparison Session 1 Bar vs Loc/L+B p=0.01). During the first trial of each session, the animal had to rely on long-term memory (2 d to 3 d between sessions) to navigate to the current GL. To minimize olfactory cues (e.g. chocolate smell and markings) the maze was cleaned with alcohol between animals, further on critical sessions (e.g. second session after a change to test for one session learning) no food was present in the maze for 60 seconds during the first trial. These probe trials were performed in session 2 and 4/5 or 6 during the Build-Up and session 2 during the Updates (Fig. 2A). Across sessions, long-term memory improved independent of the GL during the Build-Up (Fig. 2C, Session F_4,28_=4.0 p=0.01 linear contrast p=0.056, GL F_4,28_=0.4 p=0.77, interaction F_16,112_=1.1 p=0.34). In the Updates, long-term memory increased across sessions as well as differed between Update types (Session F_2,14_=3.7 p=0.053 with linear contrast p=0.009, type F_2,14_=3.7 p=0.052 with linear contrast across Bar, Loc, L+B p=0.028, interaction F_4,28_=0.58 p=0.68). Similar to the all-trials performance, in barrier Updates performance was better than in the other two types of Updates where the GL changed.

**Fig. 2.**
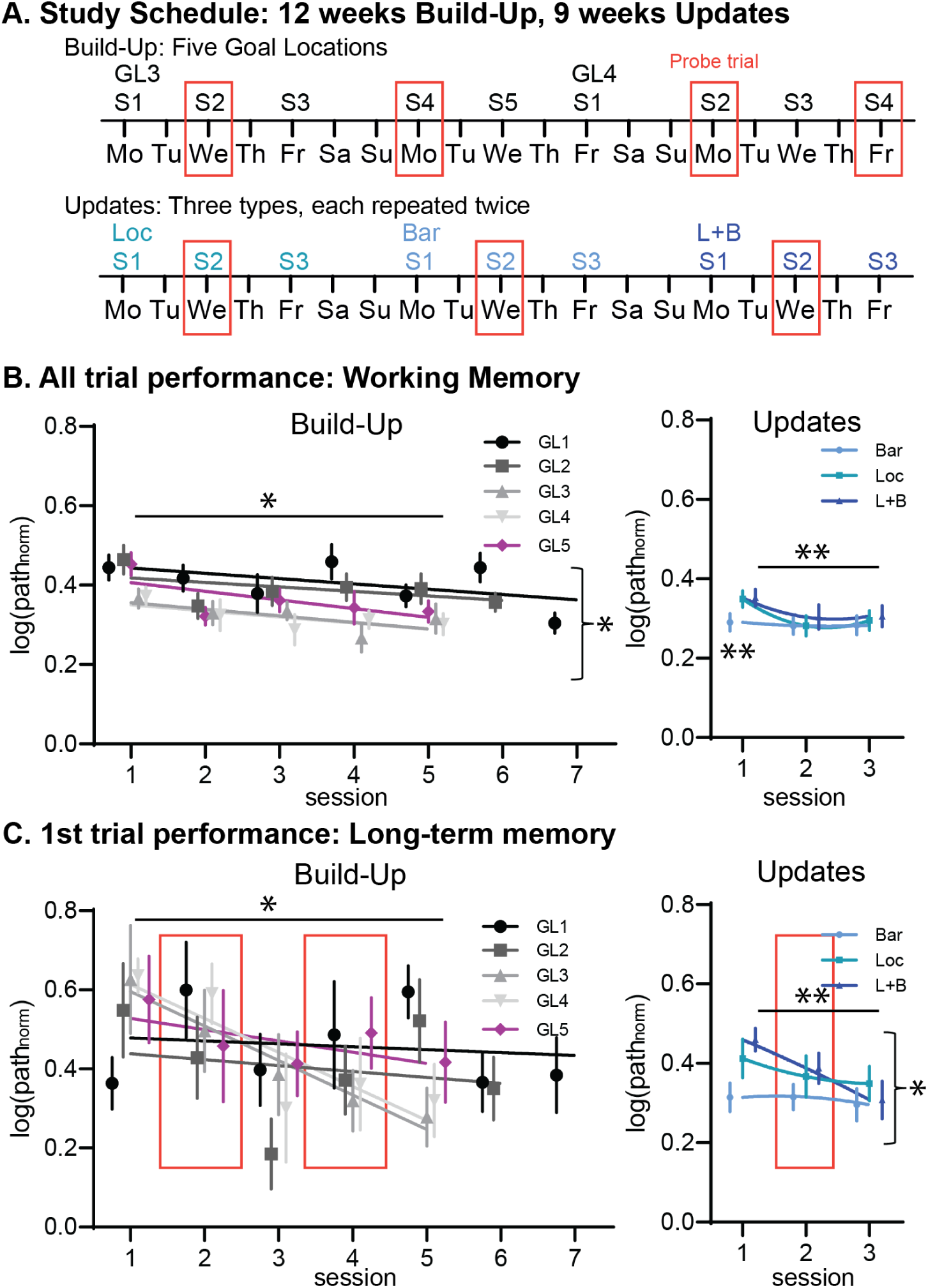
HexMaze Performance 3dw training: **A.** Shows schedule examples for the Build-up and Updates. Orange boxes indicate days with probe trials (no food for 60s of the first trial). **B**. Performance across all trials measures general working memory/navigational performance within the environment. During Build-Up there was a significant effect across session and across the five goal location (GL) switches. In contrast, during Updates, only if a location switch was involved in the update (Loc/L+B), performance was worse during the first session of the change and an improvement across sessions is visible. **C**. Performance on the first trial of each session measures the ability to remember the GL from 2 to 3 d ago. During the Build-Up long-term memory improved across sessions. During the Updates there was an improvement across sessions as well as a difference between types with larger changes in the environment (linear from Bar to both L+B) leading to worse performance. This is especially noticeable in session 1 for Loc and L+B switches where the goal is initially unknown, whereas for a Bar update only an adaption of the route is involved. Single asterisks indicate p<0.05 and double asterisk stand for p<0.01, all displayed lines in the data figures are polynomial fits.

### Time versus training

In contrast to coh 2+3, coh 4+5 (n=8) were trained only two days a week (2dw), which resulted in a shift between the training day and time alignment between both groups (Fig. 3A). As with 3dw training, 2dw lead to an improvement in all-trials measurement across sessions as well as across GLs (GL1-4, Session F_4,28_=18.3 p<0.001 linear contrast p<0.001, GL F_3,21_=4.7 p=0.011 linear contrast p=0.044); further, in contrast to the 3dw training there was a session X GL interaction (F_12,84_=2.7 p=0.004) with a faster improvement across sessions in later GLs (Fig. 3B). Long-term memory (first trial performance) improved across sessions (Session F_4,28_=12.5 p<0.001 linear contrast p=0.001) but there was no change from one GL to the next (p=0.49) as also seen with coh 2+3. Including both 2dw and 3dw in one ANOVA revealed a GL X session X training type interaction for all trials (F_12,168_=1.9 p=0.039) and for first trials a training type main effect (F_1,13_=6.7 p=0.023) as well as a marginal session X training type interaction (F_4,52_=2.4 p=0.066).

**Fig. 3.**
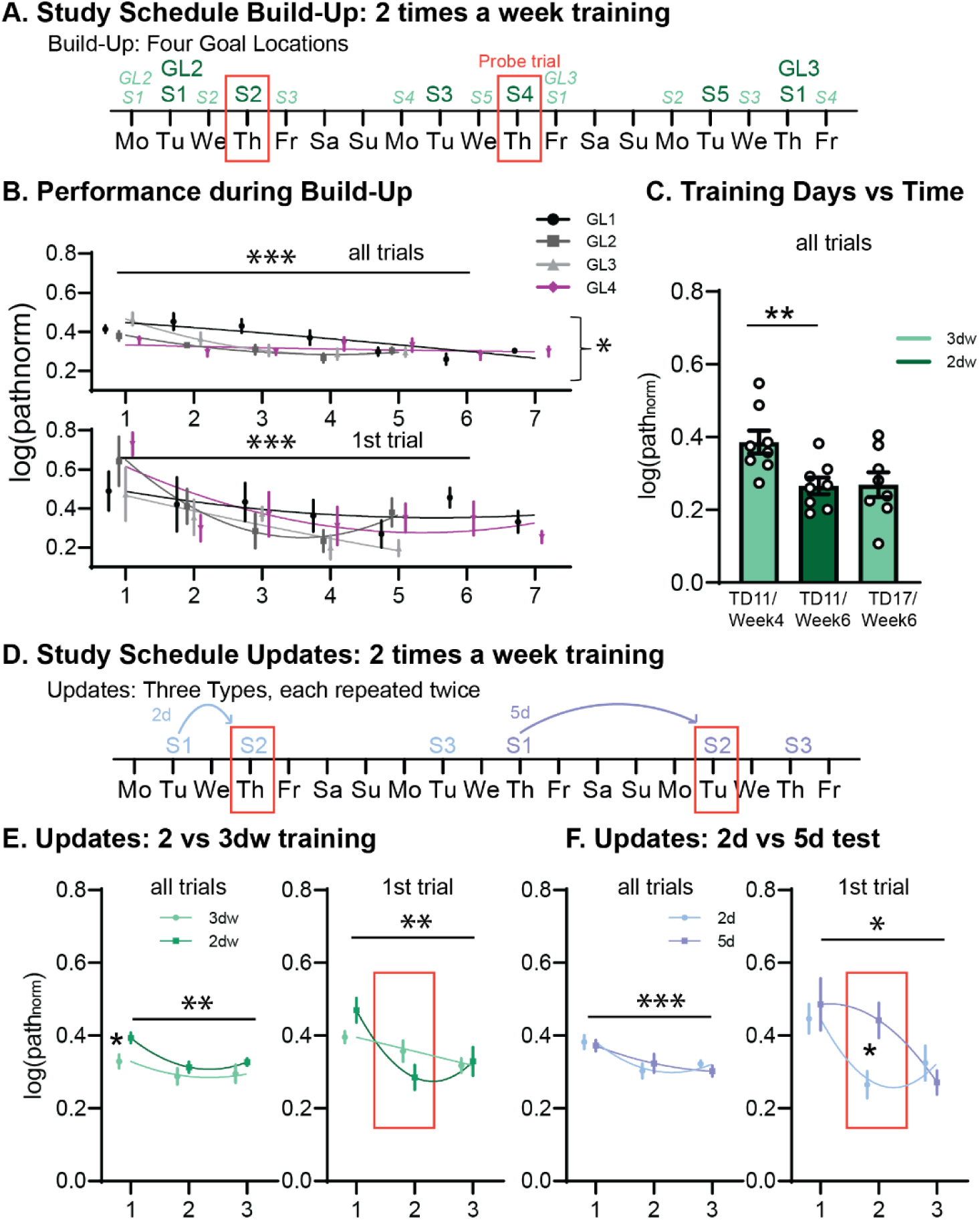
HexMaze Performance 2dw training: **A.** Shows schedule examples for the Build-up. The schedule for coh 4+5 (2dw) are shown in dark green and for coh 2+3 (3dw) in light green, illustrating the resulting shift in alignment of training days and time. Orange boxes indicate days with probe trials (food not present for 60s of the first trial). **B**. Performance across both all-trials measurement (general working memory/navigational performance within the environment) and first-trial measurement (long-term memory). We found a significant improvement in performance across sessions for both measures and additionally across GL and GL X session interaction for all trials. **C.** To compare 2dw with 3dw training, we included the corresponding training day as well as session according to time of coh 2+3 and compared these with the performance of coh 4+5. It is important to note, that the performance depended on how much time had elapsed since first exposure to the maze (weeks), not how much training the animals had received (TD is training day). **D**. Show examples from the study schedule of the Updates. With 2dw a natural alternation of two- and five-day gaps ensued during the Updates**. E.** Comparing only the 2d Updates of coh 4+5 with the Updates of coh 2+3 (also 2d gaps) showed only an Update difference during the first session**. F.** Plotted is the performance during Updates for coh 4+5 for both the two- and five-day delays. One session of training only led to significant long-term memory that lasted two not five days whereas two training sessions did indeed lead to a five-day memory persistence visible in the third session (2d condition for session 2). The single asterisk stands for p<0.05, the double asterisk for p<0.01 and the triple asterisk for p<0.001, all displayed lines in the data figures are polynomial fits.

As one of the goals was to evaluate if general performance was determined by the amount of time that had passed in contrast to how much training the animals had received, we included the same training day of coh 2+3 and coh 4+5 as well as the session of coh 2+3 that corresponded to the same week of training as coh 4+5 in a univariate analysis (F_2,21_=5.253 p=0.014, coh 2+3: training day 11, Session 4 of GL2, during week 4 and training day 17, Session 4 of GL3, during week 6, coh 4+5 training day 11, Session 4 of GL2, during week 6). Coh 4+5 performed in a similar manner to coh 2+3 when compared with how much time had elapsed, but were significantly better than coh 2+3 with the same amount of training (Fig. 3C). Thus, performance in the HexMaze was more dependent on the time period the animals had been exposed to the maze and not how much training or exposure itself was involved.

To further validate if this also applies to the previous knowledge effects, we focussed as a next step on the Updates (Fig. 3D). Both the all-trial as well as first-trial measure showed an improvement across sessions (F_2,28_=9.5 p=0.001 with linear contrast p=0.005) as well as a marginal session X training interaction (F_2,28_=3.1 p=0.06) but did not expose an effect of training amount (p=0.87, Fig. 3E). Only during the first session did coh 4+5 perform worse than coh 2+3 (p=0.01). Thus, despite the decreased amount of training, rapid updating was still possible, indicating that the creation of significant schema networks is dependent on time not training.

The 2dw training schedule also allowed us to investigate how many sessions are necessary for memory persistence as the training schedule naturally alternated with two- and five-day gaps between sessions (Fig. 3D). While one session was sufficient for the animals to remember in the first trial two days later where the food was located, this memory did not last five days (Fig. 3F). However, after two sessions of training (2d condition in Fig. 3F) the animals did remember the GL in the third session (five days after the second session, Session F_2,14_=8.1 p=0.005 with linear contrast p=0.016, interaction session X delay F_2,14_=3.6 p=0.054, delay overall p=0.34). In contrast, general navigational performance (all-trial measure) did not show a difference between the two delays (interaction p=0.24, delay p=0.9, Session F_2,14_=34.7 p<0.001 with linear contrast p<0.001).

### Previous Knowledge Effects

Different effects of previous knowledge could be observed in the resulting data, next we will focus on specific sessions and trials to highlight these effects. The simplest effect is already seen in the first GL during the Build-Up where a significant session effect indicates that each session benefits from the experience of the previous session (coh 2-5 n=16, F_6,90_=5.6 p<0.001, Fig. 4A). This simple learning effect, while often not considered as previous knowledge effect, does affect session performance and thus, must be considered even in experiments which just focus on each session individually, as seen in most electrophysiological experiments (Lopes-Dos-Santos et al., 2018; Michon et al., 2019; Roux et al., 2017).

The second previous knowledge effect can be evaluated by how well an animal can navigate within an environment and how fast this navigational capability can be adapted to a new goal as soon as it has learned a specific task. Here, this was tested at every GL switch from the beginning of the Build-Up to the end of the Updates (coh 2-5, n=16). Including the first two sessions of the first two GLs during the Build-Up as well as during the Updates (averaged across all types) revealed three distinct steps (Fig. 4B, session F_1,15_=12.6 p=0.003, GL F_2,30_=8.3 p=0.001, interaction F_2,30_=3.9; p=0.031). For the first GL, performance does not increase from the first session to the next, but as seen in Fig. 4A a performance improvement develops over seven sessions. After the first GL switch (GL1 to GL2), performance decreases to the level of performance during the first session of GL1. However, a significant improvement is exposed already for the second session of GL2 (three weeks after training start). Finally, as a third step, we find these improvements to occur in any first Update session, including additional gains in the second Update sessions (12 weeks after training start). These effects are visible across all-trial performance measurements and are likely a result of a mix of learning set effects (Harlow, 1949) as well as of an effect of increased knowledge of the maze layout. When averaging the performance across all sessions (Fig. 4C, coh 2-5, n=16), animals overall reached plateau performance already at the second GL switch during the Build-Up.

By focussing in more detail on the first and second sessions during the Updates, we can consider the amount of information animals need to incorporate during the Updates (all cohorts, n=20). We found a significant main effect of session and an interaction between session and Update types (session F_1,19_=27.5 p<0.001, type F_2,38_=2.8 p=0.072, interaction F_2,38_=7.3 p=0.002). Follow up test revealed that within session one, the amount of novel information that needs to be integrated into the existing map affects the within session online performance (just barrier, just new location or both, linear contrast p=0.03 in S1 Fig. 4D). However, this difference is eliminated by the second session, indicating that the information had been completely incorporated during the offline period.

As a final step, we tested for the schema effect – enhancement of long-term memories – by comparing the same two sessions but only including the first trial. Similar to the all-trial performance measurement, the first session performance was worse for conditions including a GL switch (Loc and L+B) as compared to just a barrier switch, but this difference disappeared by the second session (Fig. 4E, all cohorts, n=20, session F_1,19_=10.7 p=0.004, type F_2,38_=5.2 p=0.01, interaction F_2,38_=0.15 p=0.5). Finally, to investigate whether this schema effect of enhancement of long-term memory after one session learning was missing initially during Build-Up, the first trial performance during the second session of the Build-Up was compared to the first trial during the second session of the Updates (only Loc and Loc+Bar). This revealed a significant better long-term memory in the second session in the Updates in comparison to the Build-Up (Fig.4F, coh 2-5 n=16, t_15_=2.1 p=0.049).

## Discussion

In the present study we aimed at developing a new rodent task that enables the investigation of previous knowledge effects on memory encoding and consolidation. The HexMaze allows the investigation of different aspects of previous knowledge effects on performance and learning. These effects ranged from simple day-to-day performance increases, to learning-set and schema effects reflected by offline consolidation and online learning. Initial application reveals that this task can be used to test different aspects of memory while simultaneously controlling for difficulty of learning across each phase in training: from build-up of knowledge to schema updates.

Most rodent memory tasks assumes that the animal is naïve to the learned material (Arakawa & Iguchi, 2018; Dudchenko, 2004; Vogel-Ciernia & Wood, 2014). This stands in contrast to naturalistic learning, which rarely occurs in isolation of previous experience once subjects have reached adulthood. This difference in research approach may be contributing to translation failures when applying knowledge gained in animal research to human subjects (Cummings, 2018; Hartung, 2013). Previous knowledge will affect behaviour and learning (Bartlett, 1932; Harlow, 1949; Tse et al., 2007), and thus, needs to be considered when applying any particular training paradigm. To test for these effects, we utilized a large spatial environment, more naturalistic in its complexity. Mice were trained to find a food location from different starting points in a maze, thereby enforcing allocentric learning to one fixed goal location per session over two training phases: Build-Up (12 weeks) and Updates (nine weeks). During the Build-Up, the goal location was kept constant for five to seven sessions before switching to a new one, while during the Updates switches occurred every three sessions. Three different types of updates were introduced during this final phase: including barriers blocking certain paths, changing the goal location, and the inclusion of both new barrier locations and new goal locations. Across all phases, memory effects were revealed, reflected by performance increases from one session to the next (measured in the normalized path lengths). Further, four distinct previous knowledge effects modulated performance and learning.

The simplest and most obvious previous knowledge (or memory) effect is already visible in the first few sessions of the Build-Up where navigation to the invariable goal location becomes more efficient from one day to the next. This simple memory effect is what most rodent memory tasks would capture, e.g. using a radial-arm maze (Jarrard, 1995) or a watermaze, testing reference memory (Morris et al., 1982). While one could argue if this simple memory effect is a ‘previous knowledge’ effect, it is important to consider it as its simplest form: knowledge gained in previous training days effects performance the succeeding day.

The second previous knowledge effect is found when comparing the performance for the very first goal location with the performance after the first and other goal location switches. Already the second goal location exposed a significant improvement in overall navigational performance during the second session in comparison to the first, thus resulting in a different learning curve across sessions when comparing with the performance for the very first goal location. This effect is then enhanced once again during the Updates as performance improvement is already present in the first session and maintained from the first to second session as well. This is reminiscent of the learning set effect (Harlow, 1949). The results obtained in the HexMaze indicate that this learning-set effect can be expressed in three phases: (1) naïve, (2) gains after offline consolidation and (3) online as well as offline gains in the final stage. However, it remains unclear if this is the result of the animals learning the rule (there is one constant food location) or the general spatial map, but most likely it is a mixture of both.

The third previous knowledge effect is tied to the third phase of the learning set effect (corresponding to online gains) and is present across the different Update types: the amount of new information incorporated into the schema affected how rapid online learning could occur during the first session of each update. When only the general maze structure was changed (inclusion of barriers) the animals were able to rapidly adapt their routes to the goal and additional sessions were not needed to reach optimal performance. In contrast, when the goal location or both goal location and the maze structure (L+B) were manipulated, online learning was slower, resulting in a performance decrease during the first session (linear relationship with the amount of elements changed). However, offline consolidation eliminated this effect and by the second session animals performed similarly for all Update types. The fact that the degree of change in comparison to the previous learned information affected online learning and thus, perhaps initial recruitment of brain structures, could explain some differences in schema effects in previous rodent and human studies. In the original paired-associate task (Tse et al., 2007), the hippocampus was necessary during update encoding and this hippocampal involvement was also observed in a similar human schema task testing for a recently acquired, simple schema (card-location associations) (van Buuren et al., 2014). In contrast, during human schema tasks that involve long-established, real-world schemas, the hippocampus tends not to be active, and instead the prefrontal cortex directly communicates with the other cortical regions (van Kesteren, Fernandez, et al., 2010; van Kesteren, Rijpkema, et al., 2010; van Kesteren et al., 2012). It would be tempting to speculate that there may be a gradient across the complexity or extent of an existing schema, which in combination with the amount of new information overlap, results in a shift from hippocampal to cortical involvement. (1) If no schema is present, the hippocampus is necessary for weeks to months, (2) if a simple schema is present, the hippocampus is necessary for memory encoding but new information becomes more rapidly hippocampal independent and (3) if a complex schema is present, the hippocampus is not even necessary for encoding, similar to fast-mapping (Coutanche & Thompson-Schill, 2014, 2015) but also see (Cooper et al., 2019).

The fourth previous knowledge effect is reflected in long-term memory performance (first trial of each session). Initially, during the Build-Up, the animals show poor long-term (2 d to 3 d) memory after one training session to a goal location; during the Updates, the consistent development of long-term memory is accelerated and detectable in the probe trials (first trial of the second session). However, one training session only lead to a two-day and not five-day memory. For long-term memory to last five days in mice, two training sessions were required. This acceleration of consolidation has previously been linked to the schema effect (Tse et al., 2007).

Using a paired-associated task, Tse et al (2007) showed that this type of schema will allow systems consolidation (and thus hippocampal independency) to occur within days instead of weeks to months. Further, they could identify that both hippocampus and the prefrontal cortex were critical for rapid updating to occur (Bethus et al., 2010; Tse et al., 2011; Wang et al., 2012), later replicated in human subjects (van Buuren et al., 2014; van Kesteren, Fernandez, et al., 2010; van Kesteren, Rijpkema, et al., 2010; Wagner et al., 2015). Recently Ghosh and Gilboa (2014) summarized four key features of schemas: (1) an associative network structure, (2) basis on multiple episodes, (3) lack of unit detail, and (4) adaptability. The requirements are present in our task for testing spatial schema or map: the multiple extra- and intra-maze cues together with the maze-layout represent the associate network structure, training takes multiple sessions or episodes and we have shown adaptability in the Updates. Solely the lack of unit detail is hard to assess in this context. One criticism of schema tasks is that usually pre-training on the schema and the updates differ in difficulty and cognitive load because the amount of items learned in the build-up versus the update is different (Tse et al., 2007; van Buuren et al., 2014), which could account for the rapid updating effect. The advantage of our framework is that during both the Build-Up and Updates only one goal location is presented for multiple sessions, thereby keeping the task difficulty constant. Thus, we could confirm that difference in the number of presented items to be learned does not explain the rapid updating effect in schemas.

The HexMaze also revealed further interesting features of schemas in mice. Firstly, we are the first to show that the Build-Up of the schema is dependent on time but not training or experience. This was revealed by training animals either two or three times a week. When comparing these two training conditions, performance was more similar when aligned to time (weeks since start of training) then to the number of days already spent in training. Further, after the 12-week Build-Up with either 36 or 24 sessions of training, all animals showed rapid consolidation during the Updates, confirming the established schema was independent of training amount. Thus, time- and not experience-dependency indicates that build-up of a schema network requires a remodeling of the network which importantly, occurs offline and for a certain time period and cannot be facilitated by training increase. This is reminiscent of the massed vs. spaced memory effect: massed training creates a stronger initial memory, however spaced training creates a memory trace that lasts longer (da Silva et al., 2014; Nonaka et al., 2017).

Different phases in the HexMaze are optimal to apply to different types of experiments. For example, if the goal is to test classic reference memory, simply using the first seven sessions to the goal location is sufficient. In contrast, if the aim would be to measure neural correlates of navigation within an environment with many days of data for direct comparisons, training should first be to one goal location but analysis would be applied from the second session of the second goal location onwards when performance is stable over time (i.e. from ninth training day). As a third application example, the investigation of offline memory consolidation would occur during the Updates as here, each change is comparable to the next (plateau performance). One key advantage of the HexMaze to many other rodent tasks: due to the naturalistic paradigm mice rapidly habituate to the maze (two 1 h sessions of habituation with all cage mates at once primarily for stress free pick-ups with tubing) and do not require other pretraining/shaping.

We have developed a flexible rodent task in which different effects of previous knowledge on memory performance, encoding, and updating can be investigated and both offline long-term memory and online navigational performance can be evaluated separately. This task will enable future studies investigating the principles of memory updates and the involved mechanisms. While we have not yet investigated whether the schema effect of rapid systems consolidation (hippocampal independency) is present in this task as well, we do find a behavioral schema effect that is likely to be accompanied by the consolidation effect. Overall, our brains are tuned to remembering things that are new, but how novel something is will depend on our experiences (Duszkiewicz et al., 2019).

## Acknowledgments

We would like to thank Dorothy Tse and Richard Morris for fruitful discussions and CCNS in Edinburgh for lending Patrick Spooner as well as the workspace for building the maze parts. We would like to thank our master and bachelor students for helping with the rodent training: Juraj Bevandic, Natasja Rutgers, Nicole Thijsen, Tijmen van Gelder, Ece Saadet Peyaz, Gamze Naz, Eleonora Carpentiero, Femke Bakker, Luuk Meulman, Yangyang Fang, Lisann Brinckner, Christiana Vallianatou, as well as Christiana Vallianatou for generating the supplemental videos and paths.

## Author contributions

A.A. performed the experiments, created the counterbalancing sheets, analyzed the data, and supervised the students, L.B. performed the experiments, helped create the set-up and analysis pipeline, J.v.d.M, A.S., I.N.L. helped perform the experiments and supervised the students, R.E. set up the acquisition and tracking system, P.S. created the gangway system, L.G. created the HexMaze and training procedure, analyzed the data and wrote the first draft of the manuscript. All authors worked on the manuscript.

## Methods and Materials

### Subjects

Five cohorts of four male C57BL/6J mice each (Charles River Laboratories) aged two months at arrival, were group-housed in the Translational Neuroscience Unit of the Centraal Dierenlaboratorium (CDL) at Radboud University Nijmegen, Netherlands. They were kept at a 12 h light/12 h dark cycle and were before training food deprived overnight during the behavioural testing period. Weight was targeted to be at 90% to 85% of the animals’ estimated free-feeding weight. All animal protocols were approved by the Centrale Commissie Dierproeven (CCD, protocol number 2016-014-018). The first cohort (coh 1) was used to establish general maze and task parameters and entered in the current analysis only in the Updates.

### HexMaze

The HexMaze was assembled from 30 10 cm wide opaque white acrylic gangways connected by 24 equilateral triangular intersection segments, resulting in 36.3 cm distance center-to-center between intersections (Fig. 1A). Gangways were enclosed by either 7.5 cm or 15 cm tall white acrylic walls. Both local and global cues were applied to provide visual landmarks for navigation. Barriers consisted of transparent acrylic inserts tightly closing the space between walls and maze floor as well as clamped plates to prevent subjects bypassing barriers by climbing over the walls. The maze was held 70 cm above the floor to allow easy access by the experimenters.

### Video acquisition and tracking

Two USB cameras (C270, Logitech, Switzerland) were installed 2.1 m above the gangway plane with overlapping field-of-vision (FOV) to provide full coverage of the arena and reduce obstruction of vision by maze walls. Image data (15 frames/s, 800 x 600 px^2^ per camera) was acquired on a low-end consumer PC (Ubuntu 19.04, AMD Ryzen 2200G, 8 GB RAM) with a custom Python scripts (Anaconda Python 3.7, OpenCV 4.1.0) at controlled brightness and exposure levels. Images were immediately compressed and written to disk for offline analysis. In parallel, online tracking was applied for feedback to the experimenter and adjustments of the paradigm. Briefly, for each camera view a mask was generated at the beginning of the experiment based on the contrasting brightness of the maze and experimental room floor. This arena outline mask was applied to new frames and a foreground mask generated using the OpenCV MOG2 background estimation implementation (Zivkovic & van der Heijden, 2006). The resulting foreground mask was cleaned and the centroid for the largest detected foreground object in a tracking search window calculated as the putative location of the mouse in the maze. The location was smoothed over time using a Kalman filter, interpolating occasional occlusions by the maze walls and similar detection failures modes. The detected location was mapped to the closest node, and visually presented to the experimenter as well as logged for offline path analysis. Synchronization between cameras for offline analysis was enabled by presenting a blinking LED (1 Hz, 50% duty cycle) in the overlapping FOV of both cameras. Experimenters could indicate start and offset of trials using a remote presenter (R400, Logitech, Switzerland).

### Behavioural Training

After arrival and before training initiation, mice were handled in the housing room daily for 1 week (until animals freely climbed on the experimenter) and then habituated to the maze in two 1 h sessions (all four cage mates together) with intermittent handling for maze pick-ups (tubing (Gouveia & Hurst, 2017)). Mice were trained either on Mondays, Wednesdays and Fridays (coh 1-3) or Tuesday and Thursday (coh 4+5). Per training day (session) each mouse underwent 30 min of training in the maze, resulting in up to 30 trials per session, Fig. S3. The maze was cleaned with 70% ethanol between animals (later clean wipes without alcohol to avoid damaging the acrylic), and to encourage returning in the next trial, a heap of food crumbles (Coco Pops, Kellogg’s, USA) was placed at a previously determined GL, which varied for each animal. GLs were counterbalanced across animals, as well as within animals across GL switches. E.g. one out of four animals, and one out of four GL per animal would be located on the inner ring of the maze while the others were on the outer ring (to shape animal behaviour against circling behaviour). Start locations for each day were generated based on their relation to the GL and previous start locations (locations did not repeat in subsequent trials, at least 60% of the trials had only one shortest path possible, first trial was different to the last and first trial of the previous session and locations had at least two choice points distance to each other as well as the GL). On average 30 start locations were needed per day per mouse, which were generated the day before training. After the mouse reached the food and ate a reward, the animal would be manually picked up with a tube, carried around the maze to disorient the mouse, and placed at the new start location. All pick-ups in the maze were done by tubing (Gouveia & Hurst, 2017). After placing the animal at the start location, the experimenter quickly but calmly moved behind a black curtain next to the maze to not be visible to the animal during training trials.

Training consisted of two blocks: Build-Up and Updates. During probe sessions (each second session of a GL switch and additionally in Build-Up GL1: S6, GL2: S5, GL3-5 S4) there was no food in the maze for the first and ninth trial of the day and each time for the first 60 s of the trial to ensure that olfactory cues did not facilitate navigation to the GL. After 60 s food was placed in the GL while the animal was in a different part of the maze (to avoid the animal seeing the placement). All other trials of the day were run with food at the GL. Probe trials and GLs switches were initially minimized, to help shape the animal behaviour. In the first trial of the day, animals would not find food at the last presented location for both the first session of a new GL as well as probe trial days (e.g. always the second session of a new GL); thus these sessions were interleaved with normal training sessions with food present at the last known location in the first trial of the day to avoid the animals learning the rule that food is initially not provided. To measure the animals’ performance, the actual path a mouse took was divided by the shortest possible path between a given start location and the GL, resulting in the log of normalized path length (Fig. 1B) and functioning as a score value. Given a sufficient food motivation and an established knowledge-network of the maze a mouse should navigate the maze efficiently. A score of 0 indicated that the mouse chose the shortest path and navigated directly to the goal. On average, animals would improve from a 3 times to 1.5-2 times longer path length than the shortest path, corresponding to 0.4 and 0.2-3 log values. Random walks through the maze are estimated with a model to result in a 4 times longer path (0.6 in log). The normalized path length of any first trial of a session was used to measure long-term memory since training sessions were two to three days apart.

Food motivation was ensured by restricting access to food for 12 h to 24 h before training and confirmed by both the number of trials ran each day as well as the count of trials during which the animal ate food at the first encounter with the food in each trial. If animals were not sufficiently motivated, the count of both would decrease. Additionally, animals were weighted three times a week and the average weekly weight was ensured to not fall below estimated 85% free-feeding weight, which was adapted for the normal growth of each animal across time.

### Data Analysis

Normalized path length for all trials was calculated using MATLAB 2017b (MathWorks, USA). Repeated measures ANOVAs were run in SPSS Statistics 25 (IBM, USA) to determine the effect of goal location switches and session on the log normalized path length during the Build-up and across the three different types of Updates. The only between subject factor was 2dw vs. 3dw training.

**Fig S1.**
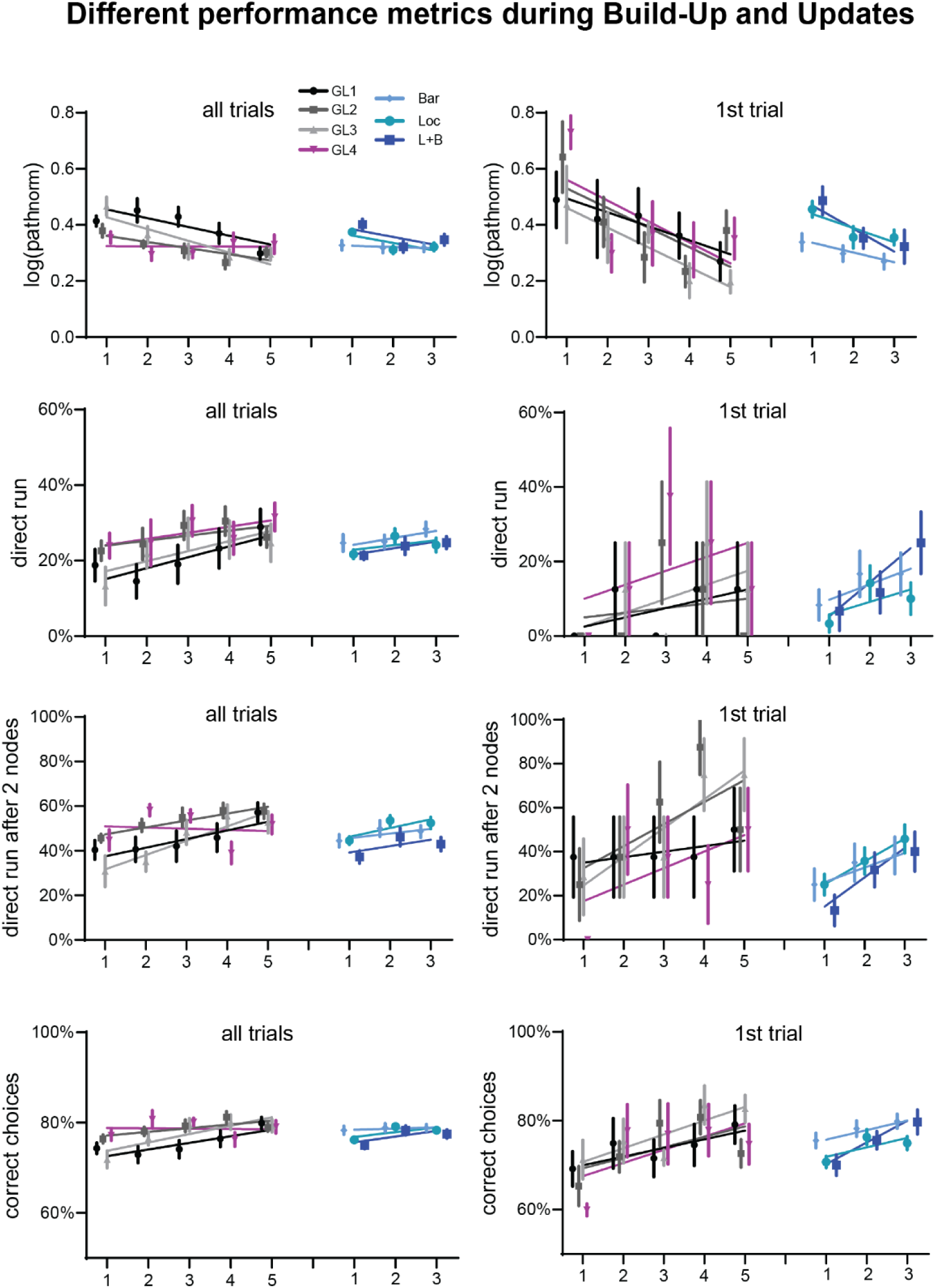
Different performance parameter Build-Up and Updates. Shown are normalized path lengths (top row, as main manuscript), percentage of trials that were a direct run (taking the shortest path, second from top), percentage of trials that were a direct run after the second node since mice often would initially run in heading direction and then stop to consider where to go (third from top). As a final parameter, we for each node the mouse traversed to the goal scored it in terms of if the choice would bring the mouse closer to the goal (correct) or not (incorrect) and created an average per trials across choices. Each left panel shows all-trials and the right panels the first trials for the Build-Up (coh 4+5) and Updates (all cohorts), all displayed lines in the data figures are polynomial fits.

**Fig S2.**
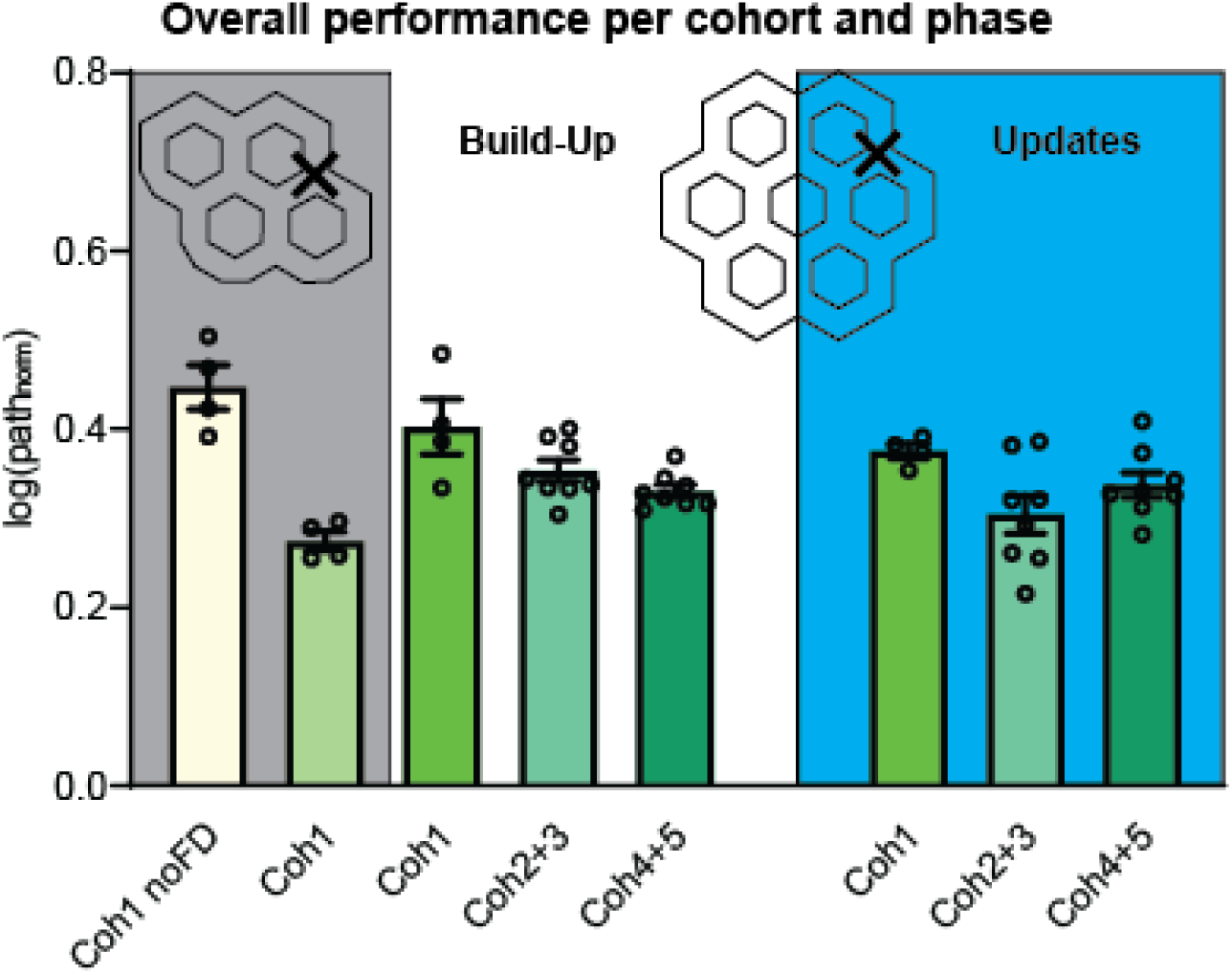
Overall performance per cohort and phase. Shown is the overall performance (all trials, sessions, goal locations) for the Build-Up (left) and Updates (right) separated per cohort. Coh 1 was initially trained on a smaller maze (left maze depiction) and not food deprived (most-left performance bar marked by the grey). However, as the animals did not perform many trials and food deprivation led to an immediate performance improvement (right bar plot marked by the grey field), later the maze was expanded while keeping the GL (as depicted on the right) to increase difficulty and later updating possibilities.

**Fig S3.**
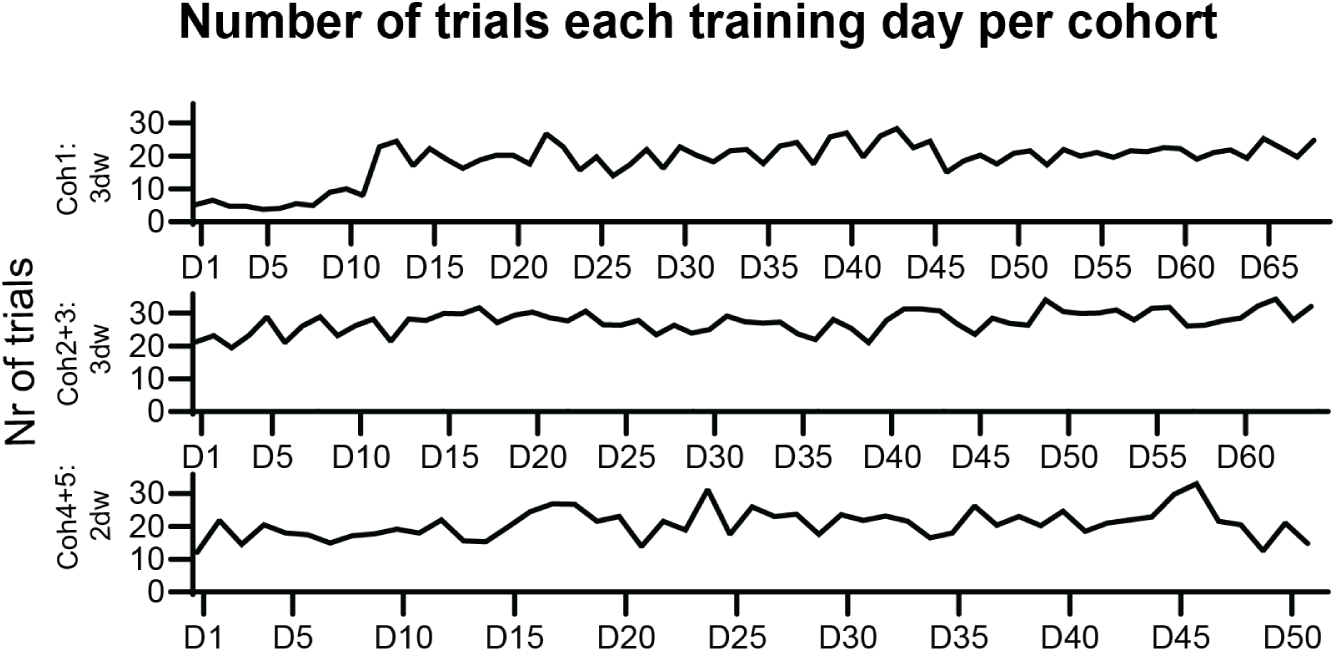
Count of trials per cohort and training day. Shown is the number of trials (averaged over cohort) that each animal ran in the 30 min session. The initial low number of trials in coh 1, were in the piloting phase when the animals were not yet food deprived.

**Video S1 Mimicking animal view in the maze**

**Video S2 Three trials of S1 during the Build-Up Mouse 12**

**Video S3 Three trials of S2 during the Build-Up Mouse 12**

**Video S4 Three trials of S1 during the Build-Up Mouse 17**

**Video S5 Three trials of S2 during the Build-Up Mouse 17**

**Video S6 Three trials of S1 during the Update Mouse 6**

**Video S7 Three trials of S2 during the Update Mouse 6**

**Video S8 Three trials of S1 during the Update Mouse 19**

**Video S9 Three trials of S2 during the Update Mouse 19**

